# The historical patterns that have shaped contemporary genetic differentiation across populations of Arctic charr in Scotland

**DOI:** 10.1101/2025.10.07.680933

**Authors:** Sam Fenton, Colin W. Bean, Kathryn R. Elmer, Colin E. Adams

## Abstract

Glacial history is an important contributor to contemporary biogeographic patterns because it caused population fragmentation and consequently diversification. The Arctic charr (*Salvelinus alpinus*) is a highly diverse non-anadromous salmonid fish species in Britain and Ireland, which likely was anadromous when it colonised around the end of the last ice age. Colonisation history of the species remains largely unexplored and so the potential impact on contemporary patterns of genetic differentiation remains unclear. To address this, we conducted a national-scale genetic study of Arctic charr using a genome-wide dataset of SNPs (24,878 SNPs and 410 individuals) and mitochondrial ND1 sequences (238 sequences). We found several mitochondrial haplotypes were shared across Britain, Ireland, and the wider Holarctic suggesting colonisation by multiple sub-lineages of the Atlantic lineage of the species. Genetic differentiation was not correlated with geographic distance among river catchments, highlighting the effect of spatial isolation and genetic drift. Several populations across different river catchments showed atypical ancestries and evidence for genetic mixing, which we speculate are due to asynchronous ice coverage and the presence of ice-dammed lakes. Our results highlight how glacial history can impact colonisation history and subsequently contemporary patterns of genetic differentiation in this widespread species.

## Introduction

Patterns of genetic differentiation between populations within a species reflect contemporary and historical processes. Differences in the environments to which populations are exposed, subsequent local adaptations, and reduction of gene flow, can lead to the accumulation of differences between populations (Orsini *et al*., 2013; Van Strien, Holderegger, & Van Heck, 2015). Many species display population level patterns of genetic differentiation that correlate with the geographic distance between populations, known as isolation-by-distance (IBD), where more geographically distant populations are more differentiated from one another. These patterns can represent differential selection on populations in different parts of the range of the species and an equilibrium between gene flow and genetic drift, often as a consequence of dispersal limitations (Hutchison & Templeton, 1999; Orsini *et al*., 2013). Other species show no relationship between population geographic distance and genetic differentiation, when levels of gene flow may reflect similarities in their environments independent of geographical distance via Isolation-by-environment (Orsini *et al*., 2013; Sexton, Hangartner, & Hoffmann, 2014). The absence of IBD occurs in panmictic species when gene flow is high and when both genetic drift and exposure to differential selection pressures throughout the range is low (Palm *et al*., 2009). Alternatively, IBD can occur when gene flow between populations is low and genetic drift is high, as is often seen in non-migratory or poorly dispersing species (Waters *et al*., 2020).

Patterns of genetic differentiation are not always a reflection of the recent contemporary processes, however, and can be influenced by a number of historical events (Rey & Turgeon, 2007; Vera-Escalona, Habit, & Ruzzante, 2015; Aguillon *et al*., 2017; Ruzzante *et al*., 2019). For example, in populations that have established relatively recently, i.e. those now inhabiting previously glaciated environments that were invaded post-glaciation, lower levels of genetic differentiation between geographically distant populations could indicate colonisation by the same ancestral population (Vera-Escalona *et al*., 2015). Similarly, presence or absence of IBD may reflect the presence of historical natural or man-made geographic barriers to gene flow (Coleman *et al*., 2018), and/or a historical change in migratory behaviour of the species (Delgado & Ruzzante, 2020).

A key driver of patterns of genetic structure for species at higher latitudes is glacial history and glaciation related processes (Hewitt, 1996; Fenton *et al*., 2023). The presence of ice sheets during glacial periods, in particular the last ice age (*ca.* 115,000 – 11,500 years ago), over large geographical scales confined many temperate species into isolated havens known as glacial refugia (Hewitt, 1999). Populations in different geographically isolated refugia had the potential to differentiate from other refuge populations over time, before separately colonising emerging new habitats in glaciated areas once they became ice-free (Finnegan *et al*., 2013; Roberts & Hamann, 2015). This has influenced species in different ways, for example, some species have ancestral lineages that stayed separate from one another when colonising (Luquet *et al*., 2019; Crotti *et al*., 2021), while others show clear points of secondary contact between the two lineages resulting in hotspots of genetic diversity (Roux *et al*., 2013; Chiocchio *et al*., 2021, 2022).

Patterns of glacial retreat during the last ice age were asynchronous, which resulted in different regions becoming ice-free at different times (Tylmann *et al*., 2022). This would have affected the timing of colonisation of previously glaciated areas and potentially which lineages may have been able to colonise. Constrained drainage of meltwater from these ice sheets in some places resulted in the formation of glacial or ice-dammed lakes which may have provided important routes for the eventual colonisation of new waterbodies by freshwater species (Fenton *et al*., 2023). Such historical lakes, that often no longer exist, may have allowed for genetic mixing across, what are now, separate river systems. This has resulted in geographic patterns of genetic connectivity that are difficult to interpret from contemporary freshwater catchments. This pattern has been shown in species like lake whitefish (*Coregonus clupeaformis*) where populations in unconnected river systems share localised ancestral histories due to colonisation through glacial lakes (Pielou, 1991; Ruzzante *et al*., 2019).

The salmonid fish Arctic charr (*Salvelinus alpinus*) is an excellent species to study the effects of historical processes on contemporary divergence, particularly in Britain and Ireland (Maitland & Adams, 2018). Arctic charr has a widespread distribution across freshwater environments in the Holarctic and displays high levels of genetic differentiation and ecomorphological variation (Jonsson & Jonsson, 2001; Adams *et al*., 2007; Elmer, 2016; Maitland & Adams, 2018). As in many other freshwater fish adaptive radiations (Taylor, Foote, & Wood, 1996; Jones *et al*., 2012; Delgado *et al*., 2019), the loss of diadromy (migration between freshwater and marine environments during the life cycle) after colonisation of emerging post-glacial aquatic systems is well documented in Arctic charr (Finstad & Hein, 2012; Jørgensen & Johnsen, 2014). While many populations in more northern latitudes still show a mixture of anadromous and non-anadromous individuals, in the southern parts of the distribution, including in Britain and Ireland, all populations are non-anadromous and no longer migrate to sea (Maitland & Adams, 2018; Salisbury et al., 2022). The species is believed to have colonised Britain and Ireland *ca.* 11 ka, towards the end of the last ice age, and these ancestral colonising populations likely were anadromous (Maitland & Adams, 2018; Fenton *et al*., 2023). As such, the contemporary, non-anadromous populations very likely arose from migratory ancestral populations, a pattern documented in other species such as alewife (*Alosa pseudoharengus*) (Palkovacs *et al*., 2008; Delgado *et al*., 2019; Delgado & Ruzzante, 2020). It is very likely therefore that for a period Arctic charr in Britain and Ireland were still anadromous and populations in different river catchments may have mixed for some time after initial colonisation. The probability of straying into a non-natal river decreases with distance, and so populations closer to each other geographically are likely to be more genetically similar (Matala *et al*., 2012; Keefer & Caudill, 2014). As such, patterns of genetic differentiation across Hydrometric Areas (defined as locations where river catchments are hydrologically connected and share the same drainage area) may reflect this period of historic anadromy, and show some correlation with geographic distance.

Another important factor that influenced patterns of genetic differentiation in Arctic charr is the number of ancestral refugia populations that colonised Britain and Ireland and the timing of those colonisations. Across its Holarctic distribution, there are believed to be five major lineages of Arctic charr (Brunner et al., 2001; Moore et al., 2015). The Atlantic lineage is prominent in Europe and holds multiple sub-lineages within it , which diverged from each other *ca.* 60 ka (Jacobsen *et al*., 2022) during the last ice age and may have colonised Britain and Ireland at the end of that ice age (Brunner *et al*., 2001; Moore *et al*., 2015). Several lakes In Scotland contain multiple distinct ecotype populations, i.e., populations that are specialised to different ecological niches within the same lake and reproductively isolated (Jonsson & Jonsson, 2001; Maitland & Adams, 2018; Fenton *et al*., 2024a). While the origins of these ecotypes in each lake are not always known, several of these ecotype divergences in Scotland are suggested to have arisen from secondary contact between different colonising populations that diverged in allopatry during the last ice age (Verspoor *et al*., 2010; Garduño-Paz *et al*., 2012; Jacobs *et al*., 2020). The colonisation history of populations across Britain and Ireland remains unknown and therefore details of how colonisation by different refugia populations impacted contemporary genetic differentiation are also unclear.

Arctic charr is believed to be one of the first colonisers of Britain and Ireland around the end of the last ice age and so would have been greatly affected by the differential timing of ice-free conditions, particularly in Scotland where the majority of contemporary populations are found (Maitland & Adams, 2018). Whilst much of Britain and Ireland was ice-free by the start of the Late Glacial Interstadial *ca.* 15 ka (Ruddiman, 2008; Clark *et al*., 2012), the northwest Highlands of Scotland remained covered in ice through this brief period of warming and until the end of subsequent Younger Dryas period (*ca.* 12.9 – 11.5 ka), also known as Loch Lomond Stadial (LLS), when glacial conditions in Britain and Ireland briefly returned before the start of the Holocene (Bickerdike *et al*., 2018). A number of contemporary populations of Arctic charr are found in regions that remained covered by ice during the LLS (Figure 1). Notably, several of these lakes (lochs Arkaig, Ericht, Lochy) contain multiple sympatric ecotype populations (Fraser, Adams, & Huntingford, 1998; Fenton *et al*., 2024b). Given that the rest of Britain and Ireland was accessible to freshwater fishes at this time, patterns of colonisation and genetic differentiation may be different for those regions that were covered by ice for longer (Cauwelier *et al*., 2018). For example, they may have been colonised by a different sub-lineage of Arctic charr to those populations located in nearby areas that were not still covered by ice. Additionally, a number of ice-dammed lakes are known to have existed in Scotland during the LLS (Sissons, 1977; Benn, 1989; Bickerdike *et al*., 2016, 2018) which may have allowed some local historical mixing between nearby populations that are now not connected.

**Figure 1:**
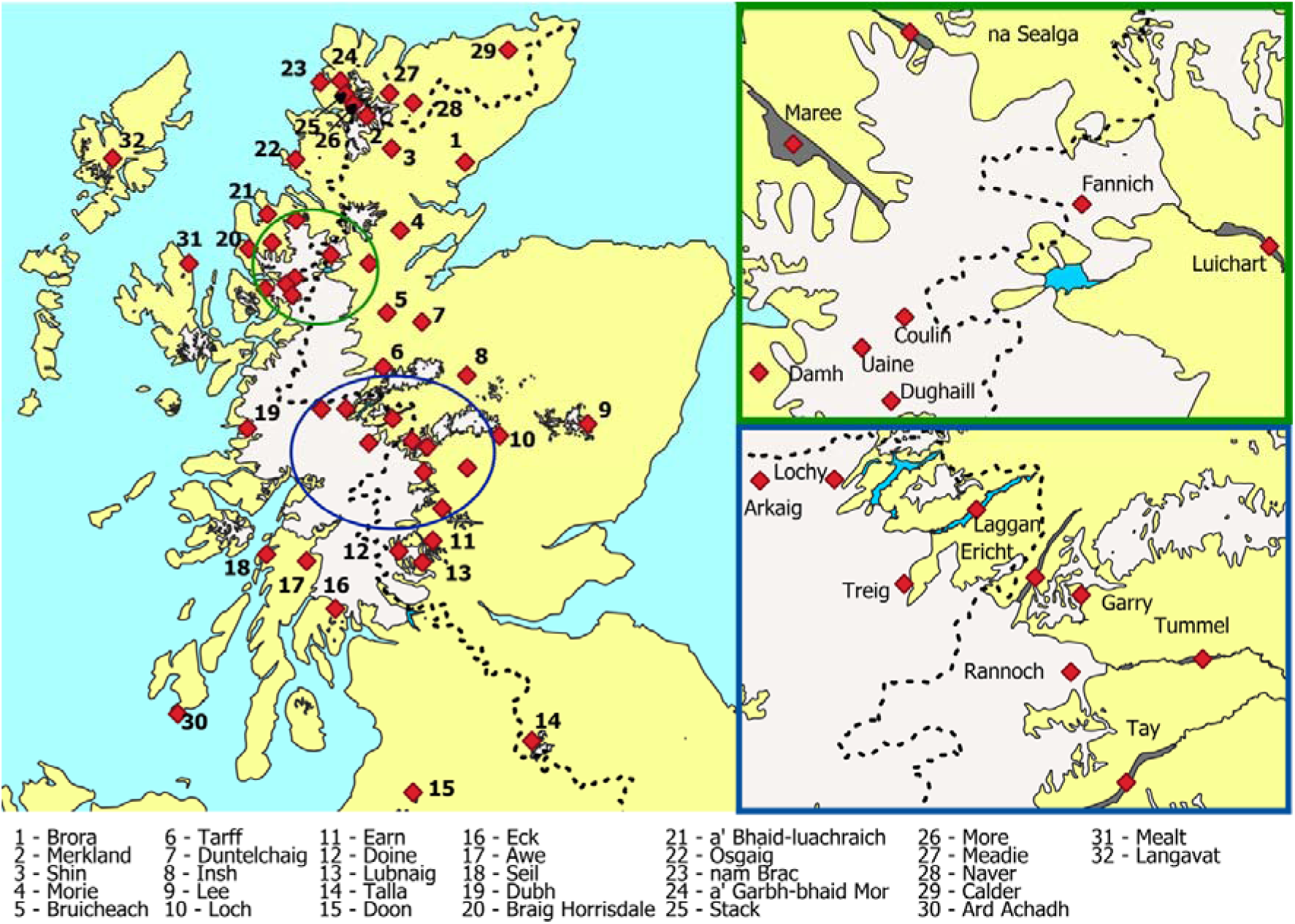
Map of ice coverage during the Loch Lomond Stadial (*ca.* 12.9-11.5 ka) in Scotland. Data on ice coverage and glacial lakes is reproduced from Bickerdike et al. 2016. Known ice cover is indicated in white and predicted ice-dammed lakes are in blue (Benn, 1989; MacLeod *et al*., 2011). Red dots indicate lakes in our dataset. Present-day lakes are shown in dark grey. The dotted black line shows the river drainage divide in Scotland, with populations to the right belonging to river system that flow out to the east coast, and populations to the left flowing out to the west coast.

In this study we evaluated the impact of historical patterns on contemporary genetic differentiation between 64 populations of the highly diverse Arctic charr found in Britain and Ireland. We hypothesised that populations in regions still covered by ice during the Loch Lomond Stadial would show distinct patterns of colonisation compared with the surrounding areas and evidence of genetic mixing across catchments (Hydrometric Areas) through known ice-dammed lakes. We first investigated whether Britain and Ireland were colonised by multiple refugia populations, using data from across the range of the species. We then explored contemporary patterns of genetic differentiation and potential migration events to identify populations with elevated genetic mixing. We complied our findings to show how patterns of ice coverage and the presence of ice-dammed lakes influenced contemporary patterns of genetic differentiation and structuring.

## Methods and Materials

### Mitochondrial ND1

We generated a dataset of mitochondrial NADH dehydrogenase 1 (ND1; 975bp in length) gene sequences for 56 populations across Britain and Ireland (N=207 individuals) (Table S1) along with two individuals from Holar (Iceland) and three from Lake Constance (Germany). ND1 was amplified from genomic DNA with PCR, using forward primer B1NDF 5’-TAAGGTGGCAGAGCCCGGTA-3’ and reverse primer B1NDR 5’-TTGAACCCCTATTAGCCACGC-3’ (Schenekar, Lerceteau-Köhler, & Weiss, 2014). We conducted PCR with 0.5 μl of forward and reverse primers at 10μM respectively. The amplification profile involved an initial step at 95°C for 5 minutes, followed 25 cycles with 55°C annealing (95°C for 30 seconds, 55°C for 90 seconds, 72°C for 30 seconds), followed by an extension at 60°C for 30 minutes. PCR product was sequenced in forward and reverse directions at Dundee Sequencing and Services (www.dnaseq.co.uk). ND1 sequences from elsewhere in the Holarctic (Canada, Alaska, Russia, Iceland, Greenland, Norway and Sweden) were drawn from Jacobs et al., (2020) or extracted from mitochondrial genomes on GenBank (Oleinik *et al*., 2020; Schroeter *et al*., 2020; Jacobsen *et al*., 2022) (Genbank accessions in Table S2). As a large number of mtDNA sequences from Greenland were available, we choose to use three representative populations for the notable lineages/sub-lineages from Jacobsen *et al*. (2021) (Arctic lineage and Atlantic sub-lineages 1 and 2), with three sequences used for each population.

All sequences were added to a multisequence alignment using *MUSCLE* in *Uniprot UGENE v45* (Okonechnikov *et al*., 2012). In total, our ND1 dataset had 238 individuals. *Salvelinus fontinalis* (Accession number: MF621738.1) and *Salvelinus namaycush* (Accession number: OM736821.1) were added as outgroups. Templeton, Crandall, and Sing’s (TCS) haplotype networks were formed using *PopArt v1.7* (Leigh & Bryant, 2015). We then ran a maximum likelihood tree using one sequence per haplotype from the haplotype network, including the outgroup species. The best nucleotide substitution model for deriving a maximum likelihood tree was selected using modeltest-NG *v.0.1.7* (Darriba *et al*., 2020). MEGA v11 was used to run the maximum likelihood tree using Tamura-Nei substitution model with 500 bootstraps (Tamura, Stecher, & Kumar, 2021).

### Genomic data

We used a ddRADseq dataset (Restrictions enzymes: MspI and PstI) previously published in Fenton et al. (2024b) (NCBI SRA BioProject PRJNA1061680, PRJNA607173), for 410 individuals covering 64 different populations (N=1-10 per population with an average of 6) across Britain and Ireland (Figure 1). Reads were mapped to the annotated *Salvelinus sp.* genome (ASM291031v2) using *bwa mem v0.7.16* (Li, 2013). RAD-loci building was done in Stacks *v2.60* (Rochette, Rivera-Colón, & Catchen, 2019) with all SNP datasets used in this study generated from this catalogue of RAD loci. Subsequent SNP filtering was performed using the *populations* module of *Stacks* and *VCFtools v0.1.16* (Danecek *et al*., 2011).

For testing genetic differentiation, SNPs were retained if they met the following criteria: present in 66% of all individuals in each population and 66% of individuals across all populations, a minimum minor allele frequency of 0.01, maximum observed heterozygosity of 0.5 with only the first SNP per locus retained and filtered to a minimum read depth of 10x and a maximum depth of 30x. For phylogenetic analyses, this dataset was further filtered to remove any SNPs with missing data along with those representing F_ST_ outliers, 95^th^ percentile for global F_ST_ scores across all individuals, or were previously identified as associated with environmental and lake variables as described in Fenton *et al*., (2025) leaving 5,949 SNPs for our 64 populations.

SNP datasets for each ecotype pair comparison were made from the *gstacks* catalogue to analyse demographic history. We did this for eight ecotype divergences from across Scotland at lochs Arkaig, Awe, Dughaill, Ericht, Lochy, na Sealga, Rannoch, and Tay (Adams *et al*., 1998; Fraser *et al*., 1998; Garduño-Paz & Adams, 2010; Hooker *et al*., 2016; Maitland & Adams, 2018; Jacobs *et al*., 2020). Due to an insufficient number of samples, the Rannoch benthivore population was excluded and so the divergence at Rannoch was run with a 2-population model (planktivore-piscivore). SNPs were retained if they met the following criteria in each comparison: present in 90% of all individuals, showed a maximum observed heterozygosity of 0.5 and one SNP per locus with no minor allele frequency filter applied to ensure rare, informative sites were retained. All SNP datasets were filtered to remove SNPs with minimum read depth < 10x or a maximum depth > 30x.

### Genetic differentiation and geographic distance

We used pairwise F_ST_ to investigate whether the extent of genetic differentiation between populations correlated with the geographic distance between them. Pairwise F_ST_ was generated using the –fstats parameter in the *populations* module of Stacks *v2.60*. Geographic distance between the lakes was calculated using *QGIS v3.26.3* to calculate the distance through river systems and the *marmap v1.06* R package for distance through the marine environment (Pante & Simon-Bouhet, 2013; QGIS Development Team, 2022). An approximate point in the middle of the lake was used as the start point for each calculation. The same point was used for lakes that contain multiple ecotypes. To properly calculate the distance across marine environments, longitude and latitude co-ordinates for a point representing the discharge point of each river to the marine environment containing one of the study populations was determined using QGIS. Marine bathymetric data for Britain and Ireland was gathered from the GEBCO database (General Bathymetric Chart of the Oceans) (https://www.gebco.net) and filtered from a minimum depth of 20 metres and a maximum depth of 200 to ensure the least-cost distance did not travel through landmasses. The least-cost distance between our marine outlet points was then calculated using the *lc.dist* function. Distance between each lake and their respective marine outlet point was calculated in *QGIS*. Marine and river distances were then combined to get the linear geographic distance between lakes in different river systems. For lakes in the same river system, the distance between the lakes along the river systems was simply calculated in *QGIS*. Populations in the same lake, i.e., multiple ecotypes, were given a geographic distance of zero. Distance values (d), genetic and geographic, were then log-transformed (D= log(d+1)) before use in Mantel tests implemented using the *vegan v2.6.2 R* package (François *et al*., 2021; Oksanen *et al*., 2022). Comparisons were split into three categories: between populations in the same lake (i.e., ecotype pairs), comparisons between lakes in the same Hydrometric Area (HA) and comparisons between lakes across Hydrometric Areas.

### Phylogenetic trees

We used *Treemix v1.13* to investigate past admixture events between populations using the same subset of no missing data SNPs (Pickrell & Pritchard, 2012). Treemix analysis was run as described in Dahms et al., (2022) and original code can be found at https://github.com/carolindahms/TreeMix. To summarise, 500 bootstraps were generated with a block size (-k) of 100 with no outgroup. A consensus tree was then formed using *PHYLIP consense v3.695* (Felsenstein, 2005). This consensus tree was then used to test for the optimal number of migration events. *Treemix* was run with the consensus tree as the starting tree (-tf) and 1-20 migration events were tested with 10 iterations each. Migration edges are added between populations in the tree to improve its likelihood. Any additional branch can be interpreted as a migration event that led to admixture. Model likelihoods for each number of migrations were compared with *OptM v0.1.*6 R package (Fitak, 2021). 30 independent runs were then performed for the optimal number of migrations using the consensus tree. The tree with the highest maximum likelihood was retained.

## Results

### Mitochondrial ND1 haplotype network

We used mtDNA sequences (ND1) from across the Holarctic range to investigate the phylogeographic history of lake-dwelling populations. From our dataset of 238 sequences, we identified 32 haplotypes, with 25 of these haplotypes found in Britain and Ireland to some degree (Figure 2). The most abundant haplotype (Brit_1) is shared across 121 of the 208 individuals and was found in 39 of 56 populations from Britain and Ireland (Figure S1). This haplotype is also found in Sweden, Germany, Iceland, Norway, and the Scoresby Sound (SCOR) Greenland populations with the sequences from Greenland, Iceland and Norway previously identified to Atlantic sub-lineage 2 (Jacobsen *et al*., 2022). Three other haplotypes found in Scotland (Scot_1, Scot_5, Scot_14) were found elsewhere in the Holarctic: in Sweden, the Disko Bay (DISK) Greenland population, and Iceland respectively with the Disko Bay population previously identified as Atlantic sub-lineage 1 and the Icelandic population of Vatnshlidarvatn previously identified as Atlantic sub-lineage 2 (Jacobsen *et al*., 2022). In our maximum likelihood tree, the sequences from Canada, Alaska and the Kapisilit river (Nuuk) in Greenland, previously identified as the Arctic lineage, split off first with the Russian sequences representing the Siberian lineage, splitting off shortly afterwards (Figure 3). This suggests that all the haplotypes found in Britain and Ireland belong to the Atlantic lineage.

**Figure 2:**
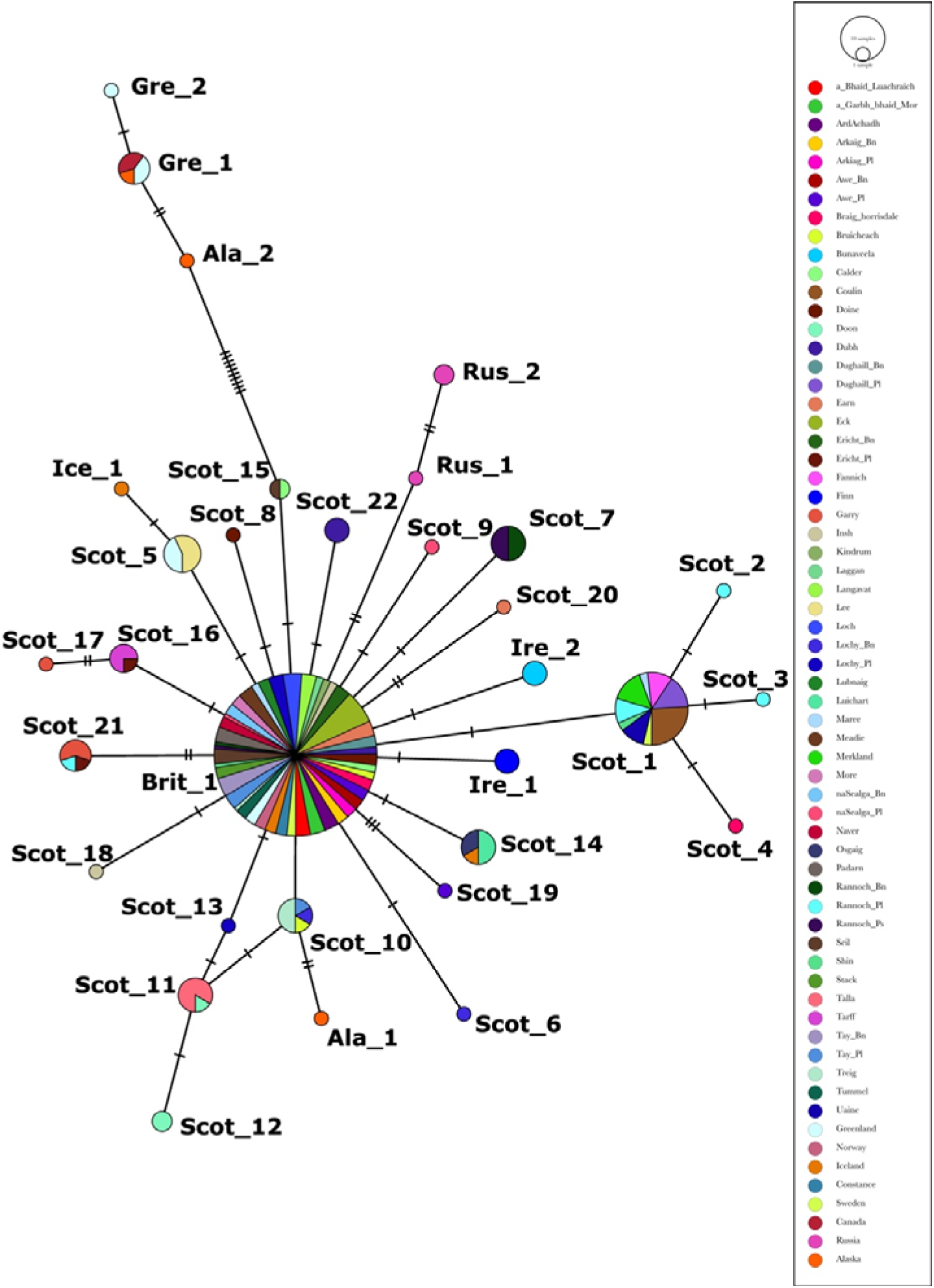
Haplotype map for our Holarctic dataset. Each point represents a population and is coloured based on haplotypes present as indicated in the legend. Panels A-F represents the different regions in our dataset: A shows North America and Greenland, B shows central Europe, C shows Iceland, D shows Ireland, E shows Wales, and F shows Scotland.

**Figure 3:**
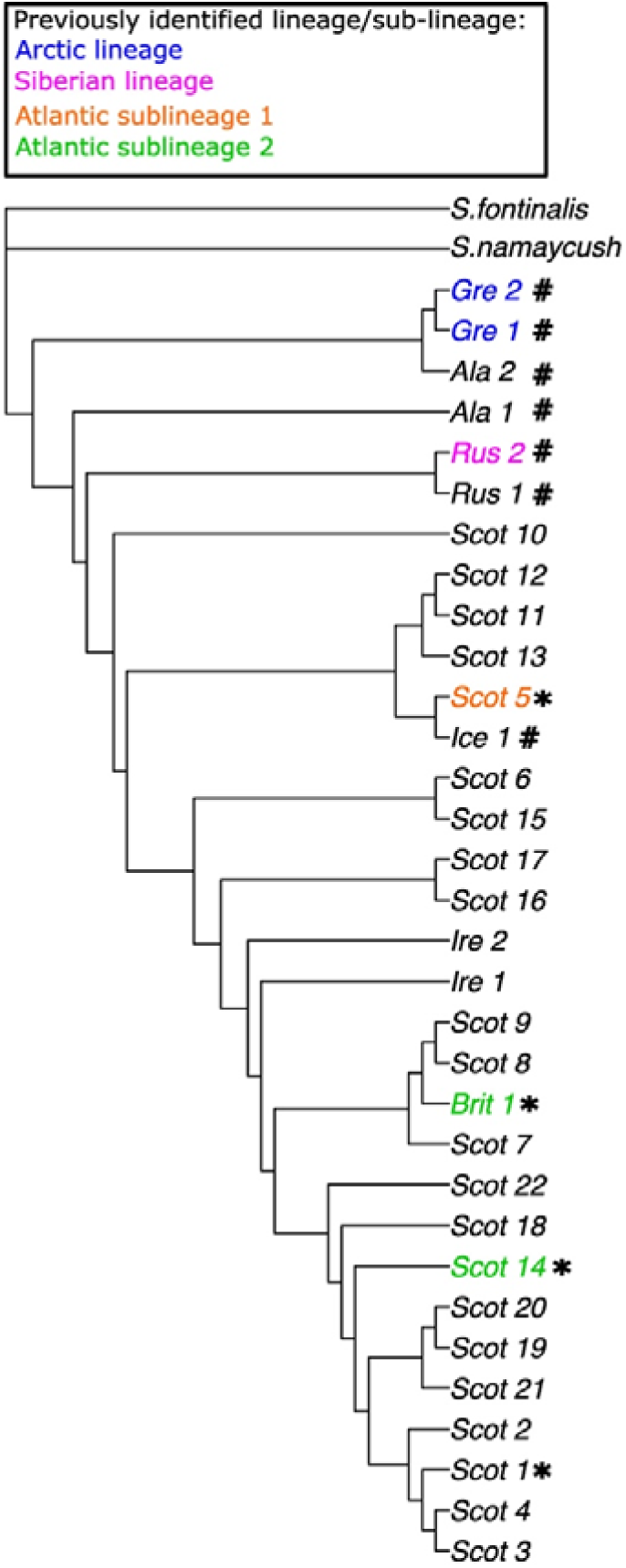
Maximum likelihood tree of haplotypes for the Holarctic dataset of mtDNA ND1. One sequence per haplotype was retained when making the tree. Haplotypes found in Britain and Ireland are named with Brit, Scot, and Ire prefix based on where they are found. Other prefixes indicate haplotypes predominantly or exclusively found in Iceland, Greenland, Alaska, and Russia. Asterisk indicates haplotypes from Britain and Ireland found elsewhere in the Holarctic. Hashtag indicates haplotypes not found in Britain and Ireland. Haplotypes belonging to sequences previously identified as a known lineage or sub-lineage are coloured to indicate their origin.

### Genetic differentiation

We used a genome wide dataset of 24,878 SNPs covering 64 populations and 410 individuals across Britain and Ireland to investigate comparisons between levels of genetic differentiation (pairwise F_ST_) and geographic distance. In total, we had 2016 pairwise comparisons which can be split into three categories: comparisons between populations in the same lake (i.e., ecotype pairs), comparisons between lakes in the same Hydrometric Area, and comparisons between lakes across Hydrometric Areas. Pairwise F_ST_ between ecotypes occupying the same lake (N=10 comparisons) ranged from 0.053 (na Sealga benthivore – na Sealga planktivore) to 0.200 (Rannoch benthivore-Rannoch planktivore) with an average pairwise F_ST_ of 0.139. Pairwise F_ST_ between lakes in the same Hydrometric Area (N=94 comparisons) ranged from 0.061 (Loch Laggan-Loch Treig) to 0.510 (Loch Loch-Loch Rannoch benthivore) with an average pairwise F_ST_ of 0.217. Pairwise F_ST_ between lakes in different Hydrometric Areas (N=1912 comparisons) ranged from 0.045 (Loch Doon-Talla Reservoir) to 0.663 (Lough Finn-Loch Mealt) with a mean pairwise F_ST_ of 0.258. Geographic distance between lakes ranged from 7.4 km between lochs Stack and More to 1359.5 km between Llyn Bodlyn in Wales and the Talla Reservoir in Scotland.

When looking across all comparisons, we saw a weak positive relationship between geographic distance and genetic differentiation (Mantel test p=0.03) with pairwise F_ST_ increasing with greater geographic distance (Figure 4A). A much stronger relationship was evident in comparisons between populations in the same Hydrometric Area (P < 0.001, adjusted R^2^ = 0.17) (Figure 4B), while no clear relationship was seen between populations across Hydrometric Areas (P = 0.011, adjusted R^2^ = 0.003) (Figure 4C).

**Figure 4:**
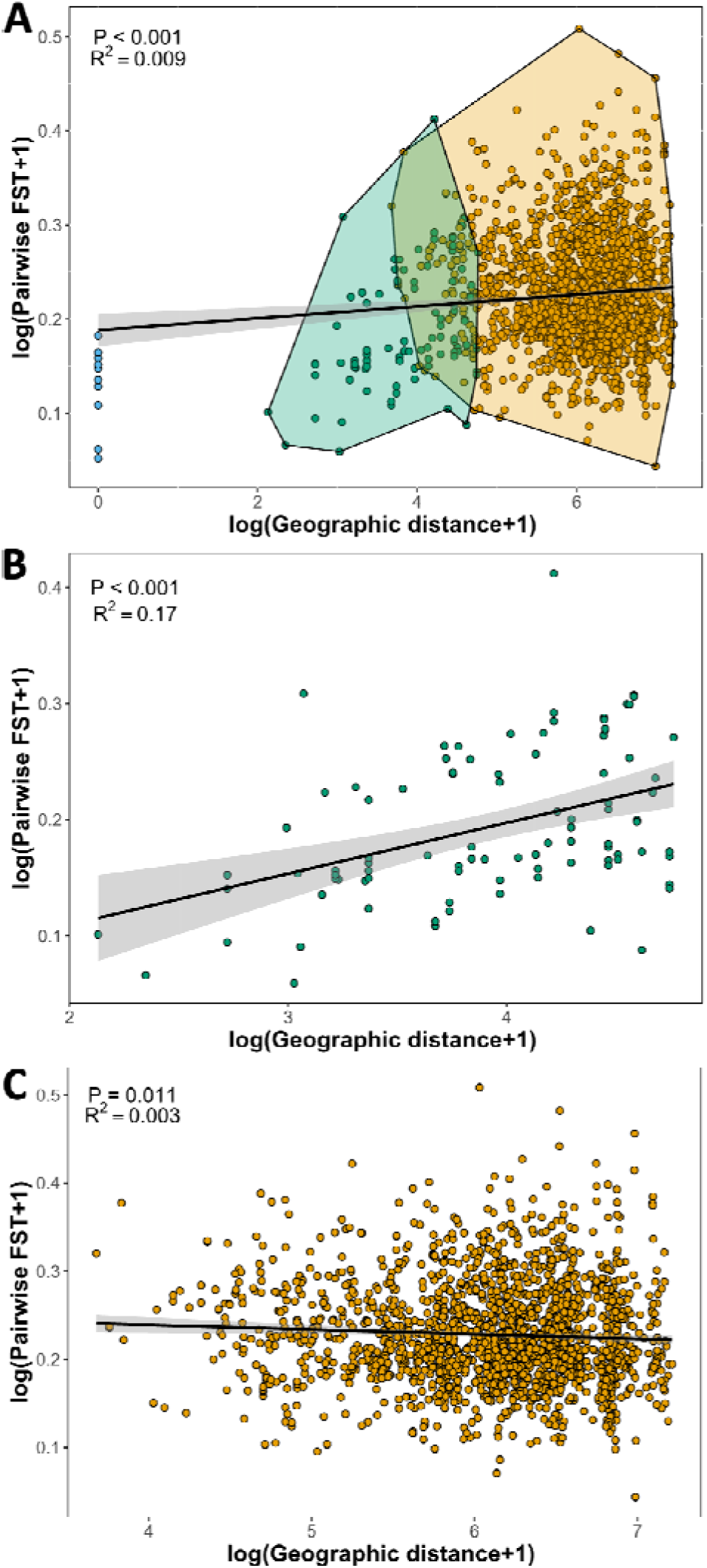
Patterns of genetic differentiation versus geographic distance between populations within Britain and Ireland (N=24,878 SNPs). Three panels show all comparisons (A), comparisons for populations within the same Hydrometric Area (B), and comparison across Hydrometric Areas (C). Black lines represent linear models with R-squared and P values displayed. Comparisons between populations in the same lake are in blue, in the same Hydrometric Area in green, and across Hydrometric Areas in orange.

### Migration events between populations

We used Treemix analysis to identify notable migration events between populations in our dataset. When we tested a different number of migration events from 1-20, and our linear models suggested no more than three notable migration events (Figure S2). These were: the migration of individuals from the Dughaill benthivore population into the Dughaill planktivore population and two migrations from populations in the Tay catchment into populations in the Lochy catchment (Loch Garry -> Loch Lochy benthivore and the Loch Ericht benthivore -> Loch Treig) (Figure 5). This shows that migration among lakes and catchments is rare and those that have occurred are concordant with the NJ tree signal of genetic-geographic discordance.

**Figure 5:**
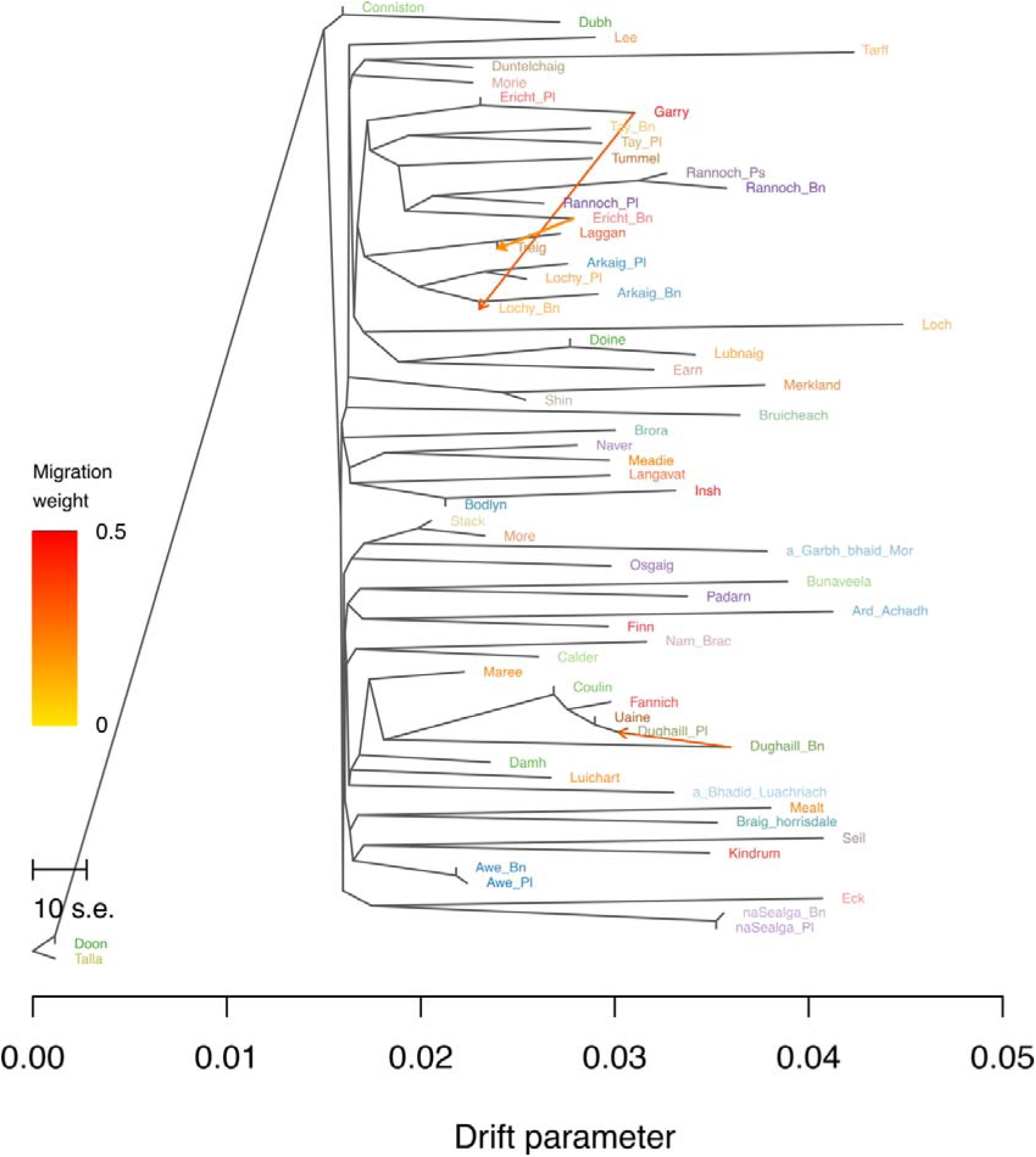
Treemix plot showing three notable migrations between populations in our dataset. Migration weight is coloured from yellow to red with increasing value.

## Discussion

Our results highlight how historical processes can provide insights that help explain contemporary patterns of genetic differentiation in Arctic charr. We identified a number of mitochondrial haplotypes found in Britain and Ireland and recorded elsewhere in the species distribution, suggesting the Isles were colonised by two known ancestral sub-lineages of the Atlantic lineage, rather than a single sub-lineage, with the potential of more sub-lineages being present (Jacobsen *et al*., 2022). Patterns of genetic structuring indicate mixing between populations in nearby river systems, likely before the anadromous phenotype was lost, and evidence of some shared colonisation histories both within and across catchments. Patterns of genetic differentiation were strongly correlated with geographic distance when looking at comparisons within the same Hydrometric Area, indicating a pattern of isolation-by-distance within catchments. Conversely, there was no relationship when looking across Hydrometric Areas, highlighting the lack of gene flow between populations in different river catchments and divergence due to drift.

We found that populations in regions still covered by ice during the likely first colonisation of Britain and Ireland by Arctic charr, the Loch Lomond Stadial (12.7 - 11.5 ka), showed genetic-geographic patterns not seen elsewhere in our dataset. This included shared mitochondrial haplotypes, evidence of genetic mixing, and low genetic differentiation across Hydrometric Areas with a known number ice-dammed lakes that existed during this time period.

### Colonisation history of populations in Britain and Ireland

Previous research has shown that Atlantic salmon (*Salmo salar*), European whitefish (*Coregonus lavaretus*), and brown trout (*Salmo trutta*) all show evidence that their distributions in Britain and Ireland comprise multiple distinct lineages or sub-lineages that diverged during the last ice age (McKeown *et al*., 2010; Finnegan *et al*., 2013; Crotti *et al*., 2020, 2021). Our findings suggest the Atlantic lineage of Arctic charr colonised Britain and Ireland, in line with other research (Brunner *et al*., 2001; Moore *et al*., 2015), and that both previously defined sub-lineages of the Atlantic lineage (sub-lineages 1 and 2) (Jacobsen *et al*., 2022) are present although to varying extents (Figure 2, Figure 3, Figure S1). No haplotypes representing the Siberian or Arctic lineage were found in our extensive sampling of Britain and Ireland, strongly suggesting that Britain and Ireland populations are solely derived from the Atlantic lineage.

Of the two defined sub-lineages of the Atlantic lineage, Atlantic sub-lineage 2 is the most prevalent in Britain and Ireland, with 41 of the 56 populations we analysed containing a haplotype shared with populations previously assigned to this sub-lineage. Such prevalent mtDNA haplotypes across large regions have been seen in other fish species such as Pacific cod (*Gadus macrocephalus*) and lionfish (*Pterois volitans*) (Toledo-Hernández *et al*., 2014; Orlova *et al*., 2019). Many of the less common Scotland-specific haplotypes also belonged to the same subclade in our phylogenetic tree as these two haplotypes (referred to here as Brit_1 and Scot_14) (Figure 3) suggesting these haplotypes also belong to the Atlantic 2 sub-lineage. Given the prevalence of the sub-lineage 2 haplotypes which were also found in Greenland, Iceland, Norway, Sweden, and Germany, this sub-lineage seems to have colonised much of the current European range for the species. Atlantic sub-lineage 1 had previously only been identified in Greenland (Jacobsen *et al*., 2022) however, one population in Britain and Ireland, Loch Lee, shared the same haplotype as was found in Greenland (Scot_5) suggesting that this sub-lineage is present in Scotland but is rare.

Additionally, we identified another haplotype (Scot_1) present both in Britain and Ireland, shared among eight populations predominately found in the northwest Highlands, and the Swedish population Sitasjaure. Given that this haplotype is shared across eight population in Scotland, is the second most abundant in our dataset, and is present elsewhere in Europe, we suggest it may represent a distinct sub-lineage. However, more data from mtDNA, such as whole mitochondrial genomes, would be valuable to infer the timing and route Arctic charr colonised Britain and Ireland from the ancestral Atlantic lineage, as has been explored in a number of other fish species (Csapó *et al*., 2020; Veríssimo *et al*., 2025). Our findings are consistent with a recently proposed scenario where the Atlantic lineage expanded from Europe, with some exemplars reaching as far as Labrador and Newfoundland (Jacobsen *et al*., 2022).

### Contemporary genetic differentiation within and across catchments

Across our dataset, we show differing relationships between the extent of genetic differentiation and geographical distance across populations (Figure 3). Comparisons between populations in the same Hydrometric Area showed a strong pattern of isolation-by-distance (Figure 3B). This pattern is in-line with evidence that migration of Arctic charr between lakes is limited (Maitland & Adams, 2018) and previous work that suggests notable gene flow only occurs between proximal populations (Fenton *et al*., 2025). Isolation-by-distance was much weaker when looking across all comparisons in the dataset (Figure 3A). Given the lack of anadromy of Arctic charr in Britain and Ireland, the need to migrate through the marine environment effectively acts as a barrier to gene flow and only generating isolation-by-distance within the same Hydrometric Areas (Figure 3B). This is similar to findings from brown trout, which only showed strong isolation-by-distance within river catchments when barriers to gene flow were removed (Griffiths *et al*., 2009) and common roach where patterns of isolation-by-distance varied by river catchment (Crookes & Shaw, 2016). Patterns of isolation-by-distance have been seen in non-anadromous Arctic charr populations in the Labrador region of Canada, although this was suggested to reflect colonisation history and high levels of admixture between colonising lineages rather than contemporary gene flow (Salisbury *et al*., 2023). We saw no relationship between geographic distance and genetic differentiation when only looking at populations in different Hydrometric Areas, populations are isolated from one another by the marine environment (Figure 3C). This reflects the lack of contemporary gene flow between these populations and highlights other processes such as genetic drift as more predominately driving genetic differentiation (Orsini *et al*., 2013; Loretán *et al*., 2020). The extent of genetic differentiation may also be driven by differences in local environment, i.e. isolation-by-environment (Orsini *et al*., 2013; Sexton *et al*., 2014) and should be explored going forward.

Previously analysis of this genomic dataset showed that broader-scale structuring, in a neighbour-joining tree, primarily split populations by east-west flowing river systems in Scotland (Fenton *et al*., 2025), which may reflect a period of historic anadromy. The finding that populations in nearby river catchments show more evidence for historic contact, in line with the probability of straying into a non-natal river decreasing with distance, has been shown in studies of Chinook salmon (*Oncorhynchus tshawytscha*) and Steelhead (*Oncorhynchus mykiss*) (Candy & Beacham, 2000; Westley et al., 2013). Given that these populations still have access to the sea and are often in close proximity to it, the loss of anadromy is likely driven either by a decrease in fitness for migrants, for example due to increase in predatory and competing species in the marine environment, or an increase in fitness resulting from not migrating (Finstad & Hein, 2012; Hogan *et al*., 2014). However, it remains unclear how long this anadromous period was, and whether the anadromous phenotype was lost for all populations around the same time.

### Glacial history shaped colonisation history and contemporary genetic patterns

Glacial history has been shown to be a key driver of patterns of genetic differentiation, due to the isolation of different refugia populations during glacial conditions and how the asynchronous patterns of ice retreat and advance influenced what regions became ice-free at what times (Fu & Wen, 2023; Fenton *et al*., 2023). Within our dataset there is evidence of how the differential patterns of ice coverage in Britain and Ireland likely influenced patterns of colonisation and gene flow. Loch Lee was the only population in our dataset to share a haplotype with the Atlantic sub-lineage 1 population (Figure 2). Notably, Loch Lee is located in an eastern region (Figure 1) that was one of the first parts of Scotland be free from ice cover, *ca.* 17 ka (Clark *et al*., 2012), a few thousand years before much of the rest of Scotland (Fenton *et al*., 2023). It has previously been theorised that sub-lineage 1 may represent an earlier colonisation event in Greenland than sub-lineage 2 (Jacobsen *et al*., 2022). Our results suggest the same maybe true in Scotland, with sub-lineage 1 potentially colonising before much of Scotland was ice-free, and as such is restricted to a few areas that were accessible during its time of colonisation.

Given that Arctic charr is estimated to have first colonised during the Loch Lomond Stadial, we propose that populations in regions still covered by ice during this time and proximal to known ice-dammed lakes would show distinct patterns of genetic differentiation. Colonisation through ice-dammed and glacial lakes has been seen in other freshwater species including lake trout (*Salvelinus namaycush*), longnose sucker (*Catostomus catostomus*), round whitefish (*Prosopium cylindraceum*), and lake chub (*Couesius plumbeus*) (Ruzzante *et al*., 2019) and can result in patterns of genetic mixing and lower genetic differentiation between populations in different river catchments (Pielou, 1991). For our dataset, there were two key areas of interest in Scotland which were both covered by ice during the LLS (Figure 1) (Bickerdike *et al*., 2016, 2018). The first of these is located in central Scotland where four populations in the ‘Lochy Hydrometric Area’ (lochs Treig, Laggan, Lochy and Arkaig), all of which are situated near a known series of ice-dammed lake that existed over Glen Spey and Glen Roy (MacLeod *et al*., 2011). The second, located in the Northwest of Scotland, is a group of populations which extend across several different Hydrometric Areas, and includes lochs Fannich, Dughaill, Coulin and Lochan Uaine, all of which surround an ice-dammed lake that was located near the village of Achnasheen (Benn, 1989).

Evidence that these populations were likely colonised through ice-dammed lakes can be seen in our study. For example, the lochs Fannich, Coulin, Dughaill planktivore and Lochan Uaine populations all consist solely of the same shared mitochondrial haplotype, despite being located in what are today three separate Hydrometric Areas (Figure 2). Notably this shared haplotype is not seen in Loch Luichart, into which Loch Fannich flows. These populations of Fannich, Coulin, Dughaill planktivore and Lochan Uaine also have low genetic differentiation to one another (pairwise F_ST_ range: 0.068 - 0.120) and previously appeared in the same section of a neighbour-joining tree (Fenton *et al*., 2025), despite relatively large geographic distances (distance range: 9.5 - 509.5 km) (Figure S3). A similar pattern has been seen in lake whitefish in North America, where populations in part of the Mackenzie drainage show more contemporary genetic similarity to populations in the Yukon drainage than to other populations found within their own catchment; these populations are close to where two proglacial lakes were found, suggesting that colonisation occurred through those lakes (Pielou, 1991). The relative proximity of these Arctic charr population to one another, all found approximately within 25 km of each other, could suggest human movement of fish between lakes (Hoffmann, 1995) however there is little evidence of this ever occurring in Arctic charr in Britain and Ireland (Maitland & Adams, 2018). As such, we propose that these genetic similarities are more likely the result of a shared colonisation history or postglacial origin, which occurred through an ice-dammed lake during the LLS.

When we looked at possible historical migration events across lakes, we found that two of them involved movement between populations from the Tay Hydrometric Area into the Lochy Hydrometric Area (Figure 5). While geographically proximal (for example lochs Ericht and Treig are approximately 20 km apart taking the geodesic distance between the two waterbodies), these two hydrometric areas flow out to the opposite sides of Scotland are approximately 958 km apart when travelling through river and marine systems. Therefore, it is unlikely these potential migrations and associated gene flow occurred via anadromous phases crossing from east to west coast across this huge geographic distance. The high genetic similarity between lochs Ericht and Treig is otherwise absent across similar geographic distances in our study. More generally, populations in the Tay and Lochy Hydrometric Areas show much lower genetic differentiation than would be expected (average pairwise F_ST_: 0.193) given the geographical distance between them (average geographic distance: 917.5km) and all populations in the west-flowing Lochy Hydrometric Area appear in the east-flowing group in the neighbour-joining plot identified in previous research on this dataset (Fenton *et al*., 2025) (Figure S3). Given that ice covered the Lochy Hydrometric Area during the LLS (MacLeod *et al*., 2011), we speculate that populations from the nearby Tay Hydrometric Area entered the ice-dammed lakes and then colonised the Lochy lakes as ice coverage retreated, similar to how a refugia population colonises the immediately around them once glacial conditions end (Herman & Searle, 2011; McDevitt et al., 2022). There are other unsampled charr populations proximal to these ice-dammed lakes that were not included in our dataset (Maitland & Adams, 2018) and warrant further investigation to test these patterns.

## Conclusions

Our results highlight the role of glacial history in shaping contemporary patterns of genetic differentiation in postglacial fishes. Arctic charr populations in lakes that were still covered by ice during the Loch Lomond Stadial showed low genetic differentiation, localised evidence of mixing between Hydrometric Areas despite a lack of anadromy, and we propose these patterns were facilitated by known ice-dammed lakes. Isolation-by-distance was strong within Hydrometric Areas, but no pattern of isolation-by-distance was found across Hydrometric Areas. Britain and Ireland were colonised by multiple refugia populations which includes the two known sub-lineages of the broader Atlantic lineage. These sub-lineages vary in their prevalence within populations and across lakes today, suggesting notably different timing of their colonisation events. These results highlight how population genetic studies benefit from considering past geography in order to interpret patterns seen in the present.

## Acknowledgments

We thank A. Niemann and A. Clark for their help in generating mitochondrial ND1 sequences that contributed to this dataset. This research was supported by NatureScot, SSE, and the School of Biodiversity, One Health & Veterinary Medicine at the University of Glasgow.

## Data accessibility

Demultiplexed ddRADseq data are available on NCBI Short Read Archive (SRA) (BioProjects PRJNA1061680, PRJNA607173). mtDNA sequence data are available on Genbank (BankIt2941812). Genetic data analysis files (vcf, mtDNA alignment) and geographic distance files, along with scripts, are permanently archived on Univ. Glasgow Enlighten (DOI XXXXXXXX [opened with acceptance])

