## Supplementary Document for "The historical patterns that have shaped contemporary genetic differentiation across populations of Arctic charr in Scotland"

**Table of Contents:**

| **Figure S1** | Page 2 |
| --- | --- |
| **Figure S2** | Page 3 |
| **Figure S3** | Page 4 |
| **Table S1** | Page 5-6 |
| **Table S2** | Page 7 |

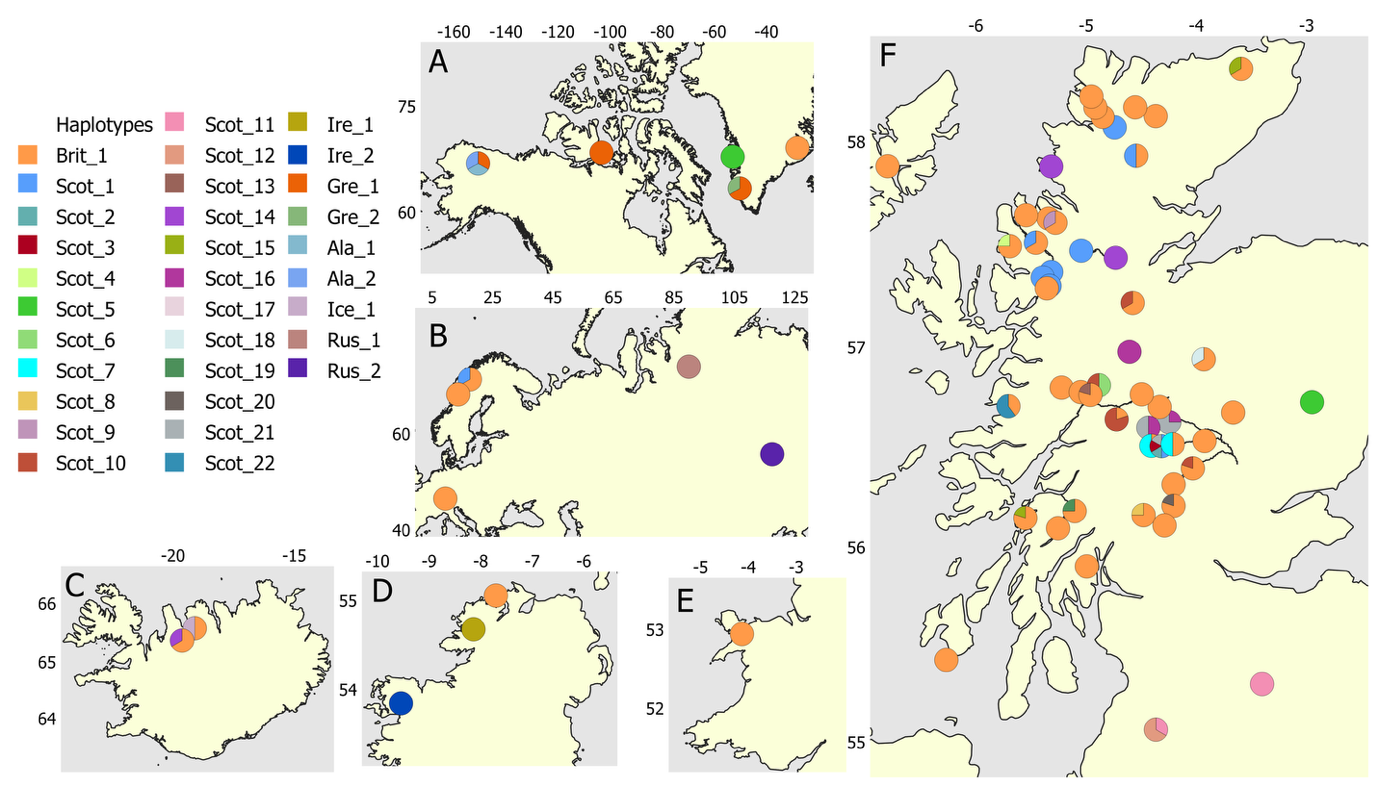
Figure S1.

Figure S1. Haplotype map for our Holarctic dataset. Each point represents a population and is coloured based on haplotypes present as indicated in the legend. Panels A-F represents the different regions in our dataset: A shows North America and Greenland, B shows central Europe, C shows Iceland, D shows Ireland, E shows Wales, and F shows Scotland.

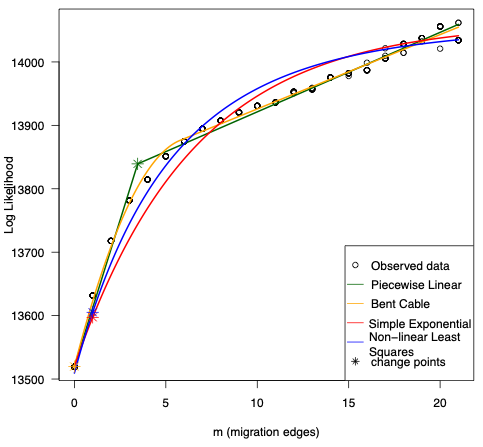
Figure S2.

Figure S2. A plot showing the most likely number of migrations from the treemix analysis. Change points indicate the most likely number of migrations for each of the fitted lines.

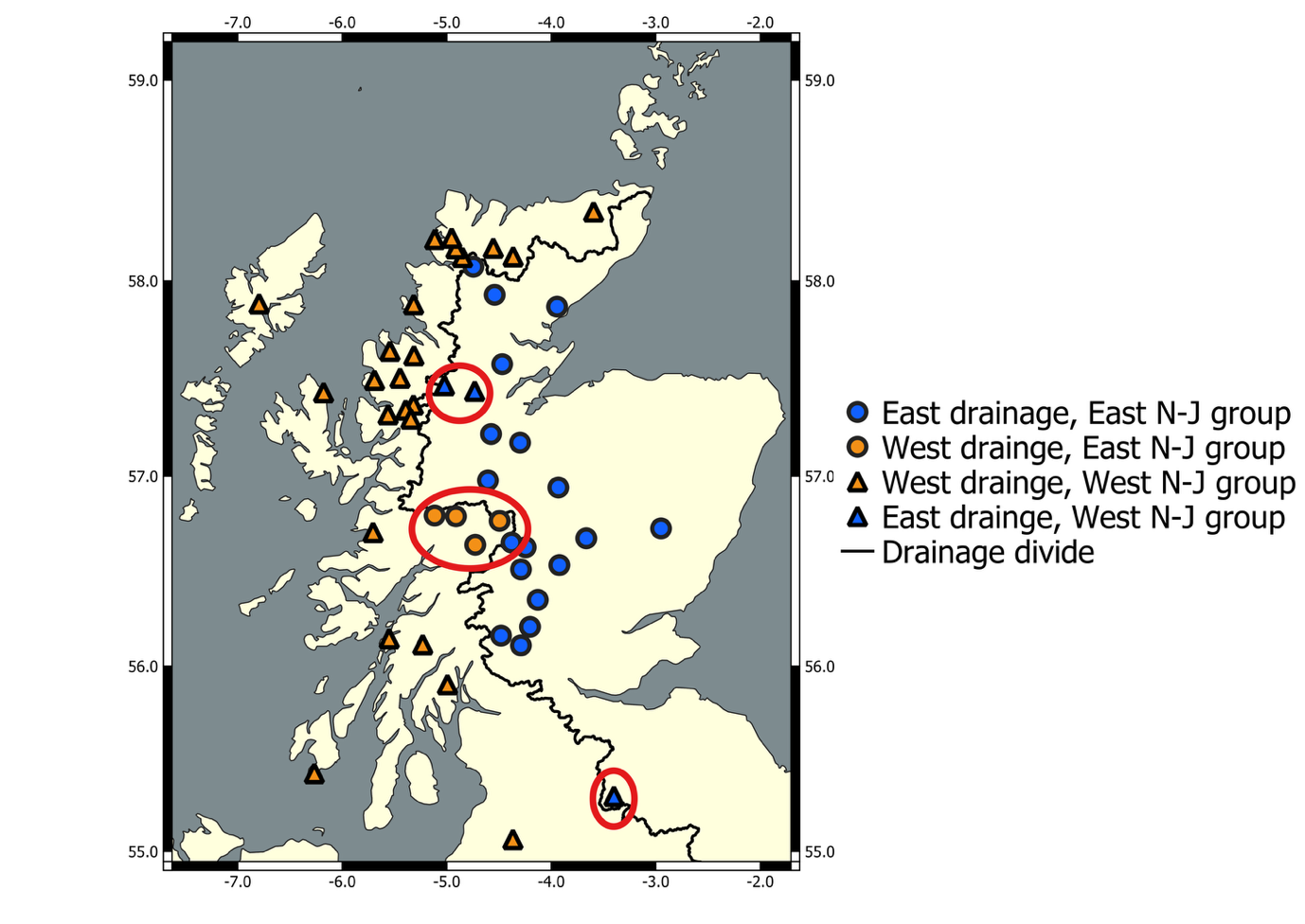
Figure S3.

Figure S3. A map of Scotland which summarises the findings from the neighbour-joining tree presented in Fenton *et al.* (2025). The map shows the correlation between placement of populations in Scotland in a neighbour-joining tree and how this correlates with which side of the central drainage divide the lake of origin is found (east vs west). Lakes are coloured by their position relative to the drainage divide (orange = west, blue = east) and shape reflects the group they belonged to in the N-J tree (triangle = west, circle = east). For the vast majority of populations, they belong to the same group in both (i.e. east drainage and east N-J group = blue circle). Red circles indicate the three groups of populations that do not follow the east-west genetic-geographic grouping. These are lochs Fannich and Luichart in the north, the Lochy Hydrometric Areas populations (Arkaig, Laggan, Lochy, and Treig) in the middle and the Talla Reservoir in the south, which was translocated there from the west-flowing Doon system (Maitland *et al.*, 2007).

Table S1.

Table S1. A summary of all populations from Britain and Ireland with have mtDNA ND1 sequences for (N=56). The number of sequences per population are indicated along with nucleotide diversity.

| Population | NmtDNA | π |
| --- | --- | --- |
| a'Bhaid-Luachriach | 4 | 0 |
| a'Garbh-bhaid Mor | 4 | 0 |
| Ard Achadh | 4 | 0 |
| Arkaig (Benthivore) | 3 | 0 |
| Arkaig (Planktivore) | 3 | 0 |
| Awe (Benthivore) | 3 | 0 |
| Awe (Planktivore) | 4 | 0.00158 |
| Braig Horrisdale | 4 | 0.00106 |
| Bruicheach | 3 | 0.00072 |
| Bunaveela | 3 | 0 |
| Calder | 3 | 0.00068 |
| Coulin | 7 | 0 |
| Doine | 4 | 0.00052 |
| Doon | 3 | 0.00068 |
| Dubh | 5 | 0.00068 |
| Dughaill (Benthivore) | 3 | 0 |
| Dughaill (Planktivore) | 4 | 0 |
| Earn | 5 | 0.00087 |
| Eck | 9 | 0 |
| Ericht (Benthivore) | 4 | 0.00068 |
| Ericht (Planktivore) | 2 | 0.00308 |
| Fannich | 3 | 0 |
| Finn | 3 | 0 |
| Garry | 4 | 0.00294 |
| Insh | 3 | 0.00068 |
| Kindrum | 2 | 0 |
| Laggan | 2 | 0 |
| Langavat | 4 | 0 |
| Lee | 4 | 0 |
| Loch | 5 | 0 |
| Lochy (Benthivore) | 2 | 0.00205 |
| Lochy (Planktivore) | 5 | 0.00042 |
| Lubnaig | 3 | 0.00072 |
| Luichart | 3 | 0 |
| Maree | 3 | 0.00072 |
| Meadie | 4 | 0 |
| Merkland | 4 | 0 |
| More | 3 | 0 |
| naSealga (Benthivore) | 2 | 0 |
| naSealga (Planktivore) | 3 | 0.00113 |
| Naver | 3 | 0 |
| Osgaig | 2 | 0 |
| Padarn | 4 | 0 |
| Rannoch (Benthivore) | 4 | 0.00054 |
| Rannoch (Planktivore) | 4 | 0.00054 |
| Rannoch (Piscivore) | 6 | 0.00177 |
| Seil | 5 | 0.00041 |
| Shin | 2 | 0.00108 |
| Stack | 3 | 0 |
| Talla | 5 | 0 |
| Tarff | 3 | 0 |
| Tay (Benthivore) | 5 | 0 |
| Tay (Planktivore) | 5 | 0.00042 |
| Treig | 4 | 0.00052 |
| Tummel | 3 | 0 |
| Uaine | 3 | 0 |

Table S2.

Table S2. Information on the ND1 sequences used from the wider Holarctic. Locality information, number of sequences used, Genbank Accession numbers and published papers the sequences were originally generated for are provided. No locality information was presented for the samples from Alaska. The samples from Holar are aquaculture samples.

| Population | Locality | Number | Accession number(s) | Paper |
| --- | --- | --- | --- | --- |
| Jayko river | Canada | 2 | MW664921.1, MW664922.1 | Unpublished NCBI submission |
| Constance | Germany | 3 | PV430342, PV430343, PV430344 | This paper |
| Disko bay (DISK) | Greenland | 3 | MT880631.1, MT880632.1, MT880633.1 | Jacobsen *et al.*, 2022 |
| Scoresby Sound (SCOR) | Greenland | 3 | MT880708.1, MT880709.1, MT880710.1 | Jacobsen *et al.*, 2022 |
| Kapisilit river (NUUK) | Greenland | 3 | MT880640.1, MT880641.1, MT880642.1 | Jacobsen *et al.*, 2022 |
| Holar | Iceland | 2 | PV430385, PV430386 | This paper |
| Vatnshlidarvatn | Iceland | 3 | MT880716.1, MT880717.1, MT880718.1 | Jacobsen *et al.*, 2022 |
| Luktvatn | Norway | 3 | MT880662.1, MT880660.1, MT880661.1 | Jacobsen *et al.*, 2022 |
| Lama | Russia | 1 | MZ571355.1 | Unpublished NCBI submission |
| Bol’shoe Leprindo | Russia | 2 |  | Jacobs *et al.*, 2020 |
| Sitasjaure | Sweden | 3 | MN957795.1, MN957796.1, MN957797.1 | Oleinik *et al.*, 2020 |
| Alaska |  | 3 | MF621740.1, MF621741.1, MF621743.1 | Schroeter *et al.*, 2020 |
